# Therapeutic Peptide SS-31 Modulates Membrane Binding and Aggregation of α-Synuclein and Restores Impaired Mitochondrial Function

**DOI:** 10.1101/2024.07.11.603085

**Authors:** Ewelina Stefaniak, Beiyuan Cui, Kexin Sun, Xucheng Yan, Xiangyu Teng, Liming Ying

## Abstract

Membrane binding and aggregation properties of α-synuclein are closely associated with Parkinson’s disease and a class of related syndromes named as synucleinopathy. This study explored the potential of SS-31 (Elamipretide), a therapeutic tetrapeptide with alternating cationic and aromatic residues and known properties of mitochondrial inner membrane binding and oxidative stress reduction, in modulating α-synuclein interaction with the lipid membranes and mitigating impairment of mitochondrial function induced by α-synuclein oligomers. It was demonstrated by both fluorescence correlation spectroscopy and fluorescence anisotropy that SS-31 displaces both wild-type and N-terminus acetylated α-synuclein from negatively charged small unilamellar vesicles in a dose-dependent manner. Thioflavin-T assay and transmission electron microscopy (TEM) showed that SS-31 inhibits membrane-induced α-synuclein aggregation and alters the morphology of α-synuclein fibrils. Moreover, Seahorse Mito Stress Test indicated that SS-31 restores impaired mitochondrial function in α-synuclein oligomer-treated neuroblastoma cells. Finally, confocal imaging revealed that SS-31 hinders cellular uptake of α-synuclein oligomers, possibly by modifying cell membrane electrostatics. These findings underscore the multifaceted protective role of SS-31 against mitochondrial dysfunction caused by α-synuclein aggregation. Consequently, SS-31 emerges as a promising therapeutic candidate to attenuate neurodegeneration pertinent to α-synuclein misfolding and aggregation. There is a good potential for further refinement of such peptide against many diseases linked to mitochondrial dysfunction and oxidative stress.

## Introduction

Substantial evidence associates α-synuclein (αSyn), a ∼14.5 kDa presynaptic protein, with the development and progression of a class of neurodegenerative disorders called synucleinopathies, including Parkinson’s disease (PD), dementia with Lewy bodies (DLB), and multiple-system atrophy (MSA).^1–4^ Intensive biophysical studies^5–7^ have indicated that αSyn is an intrinsically disordered protein (IDP), tightly coupled to the aggregation and generation of neuronal inclusions, which then accumulate in the cytoplasmic space of dopaminergic neurons, leading to mitochondrial dysfunction and cell death. Such events result in decreased dopamine levels and eventually PD symptoms.^8–11^

αSyn adopts a variety of conformational states, including a helical membrane-bound form, a partially folded intermediate state, various oligomeric species, fibrillar and amorphous aggregates.^12^ The relatively low hydrophobicity and high net charge determine the natively unfolded nature of αSyn. Thus, an increase in its hydrophobicity and/or decrease in net charge by changes in the environment surrounding the protein can induce partial folding.^13^ Upon membrane binding, disordered monomeric αSyn transits to a conformation rich in amphipathic α-helical structure promoted by seven imperfect sequence repeats of 11 amino acids with a KTKEGV consensus sequence in the region across the N-terminal and NAC domains (residues 1-90) as shown in Fig. 1A.^14^ These modular repeats provide αSyn the versatility to bind lipid membranes via multiple binding modes^15^ and exhibit different structural topologies, such as broken and fully extended α-helices.^16–18^ These helices are initiated by hydrophobic amino acid residues that interact with the fatty acyl chains of membrane lipids and by the positively charged lysine residues interacting with the negatively charged phospholipid head groups.^19^ A multilateral model was proposed to describe the lipid membrane binding structure of αSyn, where the first helix attaches the lipid membrane like an anchor while the second helix lands on the surface of the membrane.^20^ Upon binding to the lipid membrane, the aggregation of αSyn is accelerated due to its hydrophobic and aggregation-prone NAC domain which has a propensity to form β-sheet-rich oligomeric conformations.^21,22^ Such oligomeric forms of αSyn exhibit higher cellular toxicity than larger and fibrillar aggregates, which can be attributed to cytoskeletal perturbances, ER stress, mitochondrial dysfunction, increased ROS production, ion flux dysbalance, synaptotoxicity and inhibition of the cellular protein synthesis and degradation.^23–27^ Lipid binding of αSyn itself is essential for some of these observed toxic effects. Therefore, to reduce cellular toxicity induced by αSyn aggregation, one of the viable approaches would be the introduction of a compound that could disrupt αSyn/lipid binding, thus slowing down αSyn aggregation, especially at the lag phase, since the initial primary nucleation step dictates the follow-up acceleration in the aggregation process.^28,29^ In the present work, we have explored the potential of SS-31, as a membrane binding modulator, aggregation inhibitor and mitochondrial function protector against toxicity of αSyn oligomers.

**Figure 1.**
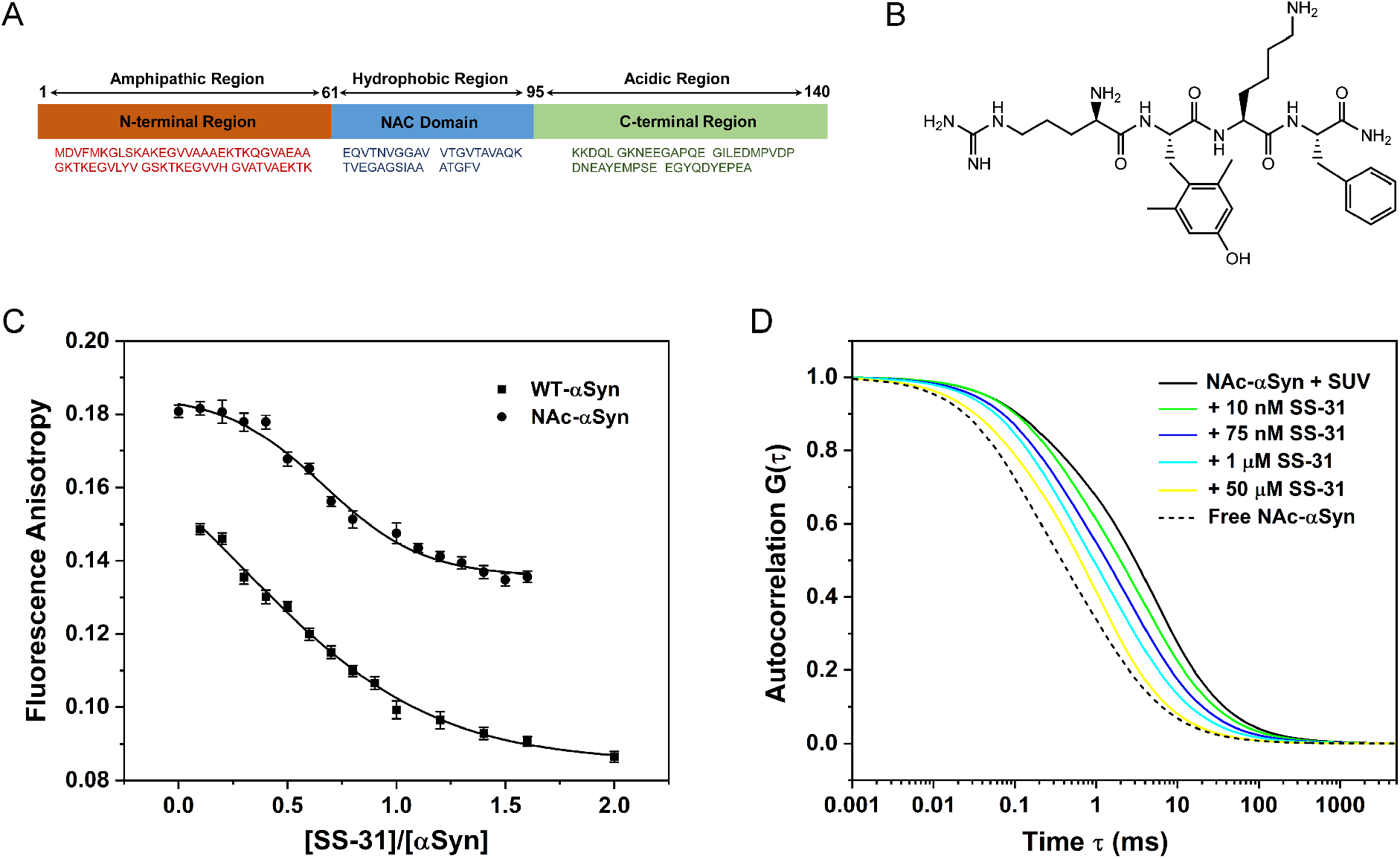
SS-31 displaces αSyn from lipid membranes. (A) Amino acid sequence of αSyn and its three distinct regions.^14^ (B) Chemical structure of SS-31. (C) Fluorescence anisotropy as a function of the concentration ratio of SS-31 to αSyn. The measurements were performed with 100 nM labelled WT-αSyn or NA-αSyn at 298 K. Error bars indicate standard deviations, n =10. (D) Fitted diffusion curves of 100 nM NAc-αSyn in the presence of 25 nM SUVs lipid upon addition of various concentrations of SS-31 and the 100 nM NAc-αSyn control (corresponding FCS curves are shown in Fig. S2 in Supplementary Information). All samples were prepared in 50 mM HEPES (pH 7.5) buffer.

SS-31 is one of the Szeto-Schiller (SS) family peptides, currently in phase II clinical trial for age-related macular degeneration^30^ and demonstrated by many preclinical studies with benefits in cardiac ageing,^31^ muscle ageing,^32,33^ atherosclerosis,^34^ ischemia,^35,36^ osteoarthritis,^37^ diabetes^38^ and glaucoma.^39^ SS-31 (also known as Bendavia, Elamipretide, and MTP-131) is a small peptide (D-Arg-dimethyl-Tyr-Lys-Phe-NH_2_, chemical structure shown in Fig. 1B) that selectively concentrates in mitochondria, suppresses and scavenges mitochondrial ROS,^40^ promotes cellular repair and restores function.^41,42^ Due to its alternating aromatic-cationic motif with a total net charge of +3 at neutral pH, this synthetic tetrapeptide binds cardiolipin (CL) enriched in the inner mitochondrial membranes via dual interactions: hydrophobic interaction with acyl chain and electrostatic interactions with anionic phosphate head groups.^43–45^ Such binding modulates the hydrophobic interaction between cytochrome c and cardiolipin^46^ and inhibits the opening of the mitochondrial permeability transition pore that forms under mitochondrial stress (e.g., traumatic brain injury, stroke, neurodegenerative diseases).^47^ Exogenously supplemented SS peptides accumulate heavily (1000-5000-fold) at the mitochondrial inner membrane (IMM) after passing through the plasma membrane in an energy-independent and non-saturable manner.^48,49^ The increase in SS-31 concentration in this region is a result of the affinity between CL and SS-31 with *n*K_D_ determined to be 2.9 µM (*n* is the number of lipid molecules bound per peptide),^45^ which could affect nearly all relevant cardiolipin-protein interactions.^50^

Mitochondria are tightly coupled to neuronal and synaptic activity, and reduced mitochondrial transport is an early event in neurodegeneration.^51–53^ Specifically, mitochondrial dysfunction is a feature observed in both sporadic and monogenic forms of PD,^54^ and notably the accumulation of αSyn within mitochondria (and subsequent formation of Lewy bodies) has been found to impair Complex I function, decrease mitochondrial membrane potential, and increase ROS generation.^55^ Preclinical studies indicated that SS-31, as one of the mitochondrion-targeting peptides,^56^ could provide therapeutic effects on neurodegeneration, e.g., protect dopaminergic neurons against Parkinsonian damage in a mouse model,^57^ reduce amyloid-β (Aβ) accumulation in a cell model of Alzheimer’s disease^58^ and enhance neural mitochondrial functions in a mouse model of cognitive deficits.^47^ A recent review systematically summarized the benefits of SS-31 therapy as well as some outstanding issues.^59^ Given that SS-31 acts on lipid membrane via the modulation of surface electrostatics of the membrane by the positive charges it carries^45^ and that the binding of αSyn to the membrane is dictated by the interaction of its positively charged resides in the N-terminus, we envisage that SS-31 might be capable of displacing αSyn on the membrane, thus inhibiting its aggregation and reducing the toxicity induced by αSyn oligomers. Herein, we report a fundamental study to uncover this unrecognized molecular mechanism that contributes to the protective role of SS-31 against αSyn oligomer induced impairment of mitochondrial function.

## Results

### SS-31 Displaces α-Synuclein from Lipid Membranes

We applied both fluorescence anisotropy and fluorescence correlation spectroscopy (FCS) to study the displacement of αSyn from lipid membranes by SS-31. To mimic lipid membranes, we employed small unilamellar vesicles (SUVs) with diameters around 50 nm, composed of DOPE (dioleoyl-phosphoethanolamine), DOPS (dioleoyl-phosphatidy-L-serine) and DOPC (dioleoyl-phosphatidylcholine) (5:3:2 w/w). The combination of these lipids is a good representation of the lipid composition of the synaptic vesicles and was commonly used to mimic them.^60,61^ Both wild-type αSyn (WT-αSyn) and N-terminal acetylated αSyn (NAc-αSyn) were used since analysis of the forms of synuclein present in LBs from DLB patients, revealed that N-terminal acetylation was a common posttranslational modification of this protein.^62^ Both forms of αSyn were also introduced with a cysteine mutation at G7 position and were labelled by Alexa 488 dye (see Materials and Methods in Supplementary Information). The process of SS-31 replacing αSyn from vesicles was initially investigated by fluorescence anisotropy. First, the binding between αSyn and SUVs was studied by titration of 100 nM labelled WT-αSyn and NAc-αSyn with the lipid vesicles (Fig. S1). Equilibrium dissociation constants (*K*_d_) derived for WT-αSyn and NAc-αSyn were 16 ± 0.5 µM and 9.5 ± 2.9 µM, respectively. NAc-αSyn exhibited a larger anisotropy value under the same condition, possibly because the 1-14 amino acid residues of N-terminal acetylated protein can be inserted into the membrane ∼2 Å deeper than WT-αSyn,^20^ further limiting the rotation of the fluorophore covalently attached to the cysteine mutation upon membrane binding. SS-31 titration was then performed after the completion of αSyn binding to the membranes indicated by the anisotropy values reaching the plateau. Fig. 1C shows the fluorescence anisotropy curves as well as fits to a dose-response equation (see SI for details) for membrane-bound WT-αSyn and NAc-αSyn with increasing SS-31/αSyn ratios. The two derived half-maximal effective concentrations (EC50) were identical, 66.5 ±1.9 µM and 67.0 ± 3.3 µM for WT-αSyn and NAc-αSyn, respectively, indicating that the SS-31 followed the same mechanism competing with wild type and acetylated αSyn. However, at high SS-31 concentration (∼150 μM) where the competition process almost reached the end, the anisotropy of the NAc-αSyn/SUV sample failed to return to the original value corresponding to that of the free protein, suggesting that acetylated αSyn not only has a higher membrane affinity but also a fraction of membrane-bound population might need to overcome a higher energy barrier (kinetically slow) to be detached from the membranes once their N-terminal residues are buried deep into the membrane.

To further consolidate the evidence for the competitive replacement between αSyn and SS-31 on SUVs, an independent method, fluorescence correlation spectroscopy (FCS), was employed. At first, the SUV/WT-αSyn conjugates were pre-formed by mixing 50 nM SUVs (total lipid 1 mg/ml) with 200 nM labelled αSyns at equal volume and incubated at room temperature for 5 minutes before measurement to ensure effective binding. Under such conditions, the protein molecules were membrane bound but the binding was not saturated since each SUV can bind on average about 45 labelled αSyn molecules.^63^ Then SS-31 was titrated into the sample to gradually replace αSyn from the SUVs. Normalized autocorrelation curves for the SUV bound WT-αSyn and NAc-αSyn treated by SS-31 together with the protein alone as control are presented in Fig. S2A&2B in Supplementary Information. Autocorrelation curves were fitted to a two-species diffusion model except for the protein alone curve where a simple one species diffusion model was used. Fitted diffusion curves are shown in Fig. S2C for WT-αSyn and Fig. 1D for NAc-αSyn, respectively. A reduction trend of diffusion time can be clearly observed from both the original autocorrelation curves and the fitted diffusion curves as SUV bound αSyn molecules were gradually replaced by SS-31, in good agreement with the fluorescence anisotropy measurement.

Taken together, the changes observed in fluorescence anisotropy and diffusion time from both assays unequivocally confirm the displacement of both forms of αSyn from negatively charged lipid membranes by positively charged SS-31.

### SS-31 Modulates Membrane-Induced Aggregation of αSyn

Taking consideration of the observation of the displacement of αSyn from lipid membranes by SS-31, we asked if SS-31 would make an impact on lipid membrane-promoted αSyn aggregation. We investigated such an effect using DMPS (1,2-dimyristoyl-sn-glycero-3-phospho-L-serine) as a model membrane system since studies have shown that DMPS is a vital component within dopaminergic synaptic vesicles and DMPS vesicles are capable of significantly promoting αSyn amyloid formation due to the negative charges on their surface.^28,64–66^ We carried out Thioflavin-T (ThT) assay with samples containing αSyn alone, αSyn with DMPS lipid, and αSyn with DMPS lipid in the presence of different concentrations of SS-31. As shown in Fig. 2A, fluorescence time profiles for WT-αSyn with DMPS lipid evolved significantly with increasing concentration of SS-31. The elongation phase of the aggregation corresponding to β-sheet formation and protofibrils growth, was substantially prolonged for samples in the presence of SS-31. Moreover, as the concentration of SS-31 increased, the elongation phase was delayed. The amount of fibrils formed was generally lower in comparison to the case without SS-31. These results suggest that SS-31 must have exerted an inhibitory effect on lipid-induced aggregation of αSyn in a dose-dependent manner. SS-31 progressively coated the surface of the membranes thanks to its efficient binding to the anionic head groups of the DMPS lipids, resulting in a reduction of available binding sites and electrostatic forces required for αSyn-DMPS binding, hence hindering membrane-promoted αSyn aggregation.

**Figure 2.**
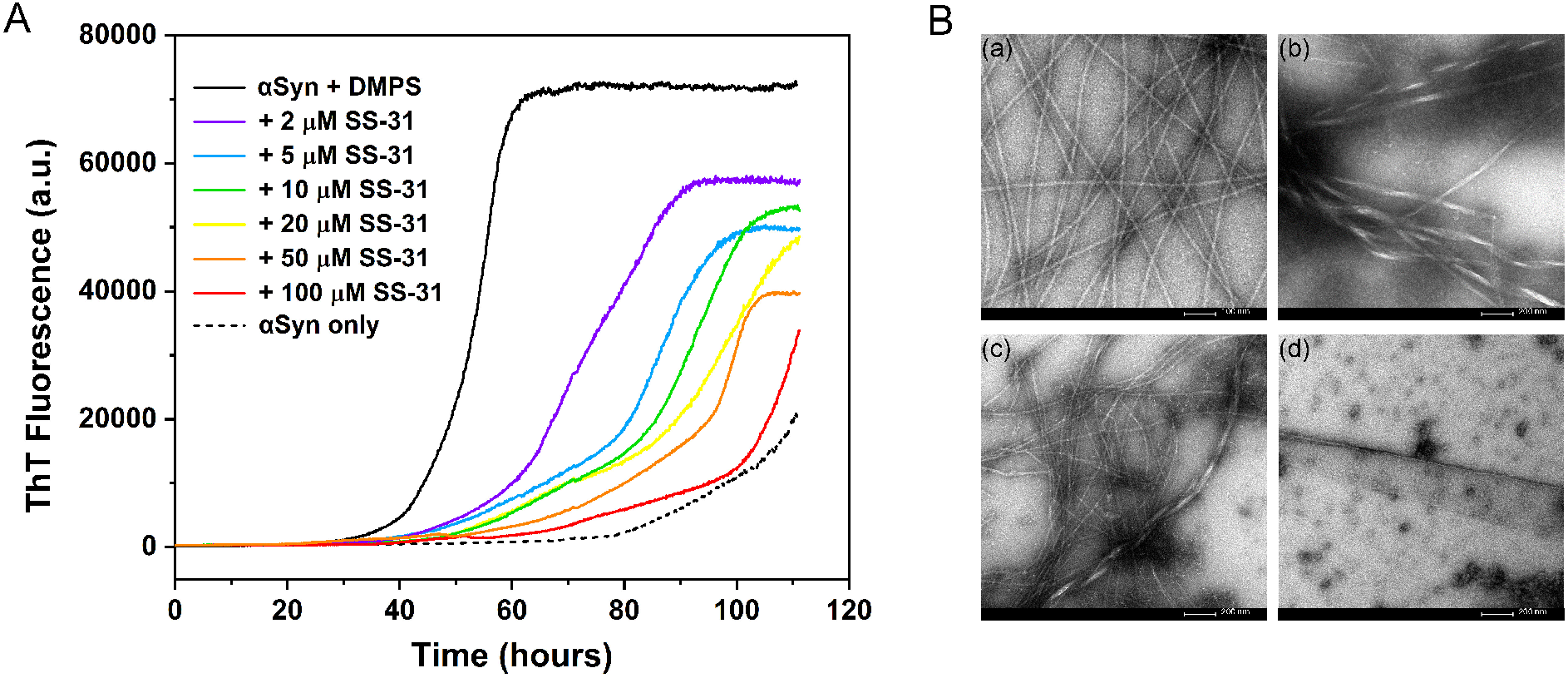
SS-31 inhibits lipid-induced aggregation of αSyn. (A) Representative ThT fluorescence time profiles for 100 µM αSyn incubated with 100 µM DMPS lipid in the presence of increasing concentration of SS-31. All measurements were run in triplicates and acquired under quiescent conditions at 37 °C (See Materials and Methods in Supplementary Information). (B) Transmission electron microscopy (TEM) images of ThT assay samples. (a) free αSyn; (b) αSyn + DMPS; (c) αSyn + DMPS + 10 µM SS-31; and (d) αSyn + DMPS + 100 µM SS-31.

Next, we investigated the influence of SS-31 on the morphology of the αSyn aggregates formed during ThT assay using transmission electron microscopy (TEM). Significant differences were found between the samples in the presence and absence of SS-31. As shown in Fig. 2B, αSyn alone formed thin and straight fibrils, while aggregates and twisted fibril bundles were observed in the presence of 100 µM DMPS lipid, consistent with previous reports.^67,68^ These large macromolecular structures formed in the presence of lipid vesicles are so-called large helical assemblies^65^ or proto-fibrils.^64^ The formation of the extended helical structure confirms the modulation of misfolding and aggregation of αSyn by lipid membrane binding.^69^ However, in the presence of both 10 µM SS-31 and 100 µM DMPS, the assemblies were mostly composed of straight fibrils or fragmented fibrillar species, but minor species, twisted and curlier fibrils, were also present. Thus, SS-31 may have helped αSyn fibrils retain their normal morphology under certain conditions. Nevertheless, the fraction of such fibrils was significantly reduced upon further increase of SS-31 concentration. 100 µM SS-31 inhibited αSyn fibril formation, probably by stabilizing off-pathway assemblies, e.g. amorphous aggregates. Collectively, observations from TEM confirm that SS-31 can induce significant and concentration-dependent inhibition of αSyn fibril formation.

### SS-31 Reverses Mitochondrial Dysfunction Induced by α-Synuclein Oligomers

As reported previously, SS-31 replaces αSyn from cytomembrane and exhibits potential to modulate mitochondrial oxidative stress in various disease models.^31,70^ Prolonged treatment with SS-31 improves function in aged mitochondria by increasing mitochondrial ATP production and inhibiting mitochondrial permeability transition pore (mPTP) formation, as well as reducing oxidative stress.^33,44,71^ To evaluate whether SS-31 exposure could improve the robustness of healthy cells and rescue mitochondrial respiratory function impaired by αSyn oligomer, we conducted a mitochondrial stress test using the Seahorse XFe96 analyzer to evaluate the bioenergetics profile of neuroblastoma cells treated with αSyn oligomer and SS-31.

As shown in Fig. 3, there was a reduction of basal respiration of SHSY-5Y cells incubated with 0.3 μM αSyn oligomers compared to the control but SS-31 treatment induced an increase in basal respiration. The maximal respiration attained by adding the uncoupling agent, carbonyl cyanide-4 (trifluoromethoxy) phenylhydrazone (FCCP), which disrupts mitochondrial potential and stimulates the respiratory chain to operate at maximum capacity, also displayed an upward trend as the concentration of SS-31 increased. At 10 μM SS-31 concentration, both the basal and maximal respiration were restored to the levels higher than that of the untreated cells. A reduction of respiration was also observed upon further increase of SS-31 concentration. Taken together, these results indicate that SS-31 can reverse mitochondrial dysfunction induced by α-synuclein oligomers.

**Figure 3.**
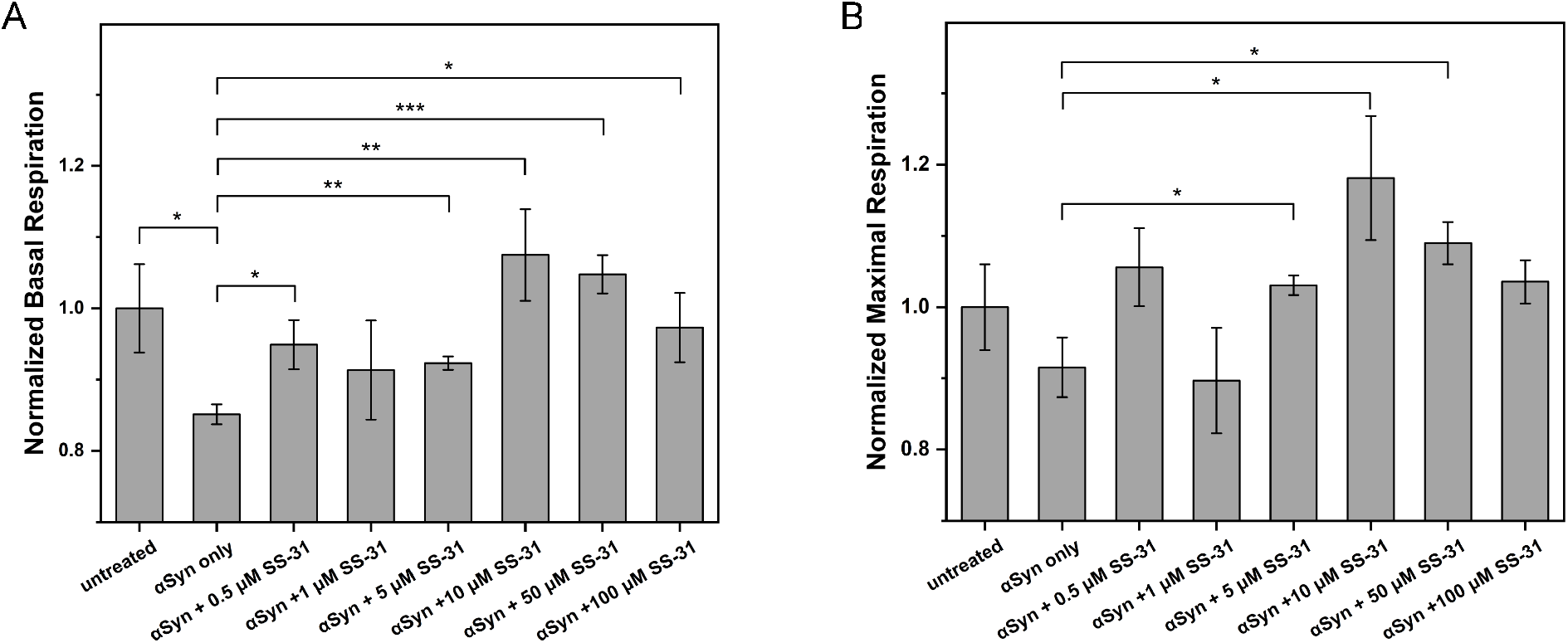
Mitochondrial respiration function of SHSY-5Y cells incubated with 0.3 μM of αSyn oligomers and treated with SS-31 (Seeding density: 20,000 cells/well). (A) Basal respiration. (B) Maximal respiration. Oxygen consumption rates were normalized to the control (untreated cells). Statistical analysis was performed with student’s T-test (*: P<0.05; **: P<0.01; ***: P<0.001). Error bars indicate standard errors of the means (n=4).

### SS-31 Hinders Cellular Uptake of αSyn

Finally, we examined the effect of SS-31 on cellular αSyn uptake by confocal microscopy. As shown in Fig. 4, αSyn oligomers were internalized to BE(2)-M17 cells but their amount in the cells was lowered after SS-31 treatment. The reduction was also dose-dependent (10 µM vs 100 µM), indicating that SS-31 restricted cellular αSyn uptake, possibly via the modulation of cell membrane electrostatics, making it less negatively charged, thus preventing the entry of αSyn. However, further escalating SS-31 concentration beyond 100 μM resulted in an increase of apoptotic cells (data not shown), perhaps due to the overload of SS-31 on cell membranes, especially in the inner membrane of mitochondria, reducing cell viability, consistent with the reduction of basal and maximal respiration observed at higher SS-31 concentration.

**Figure 4.**
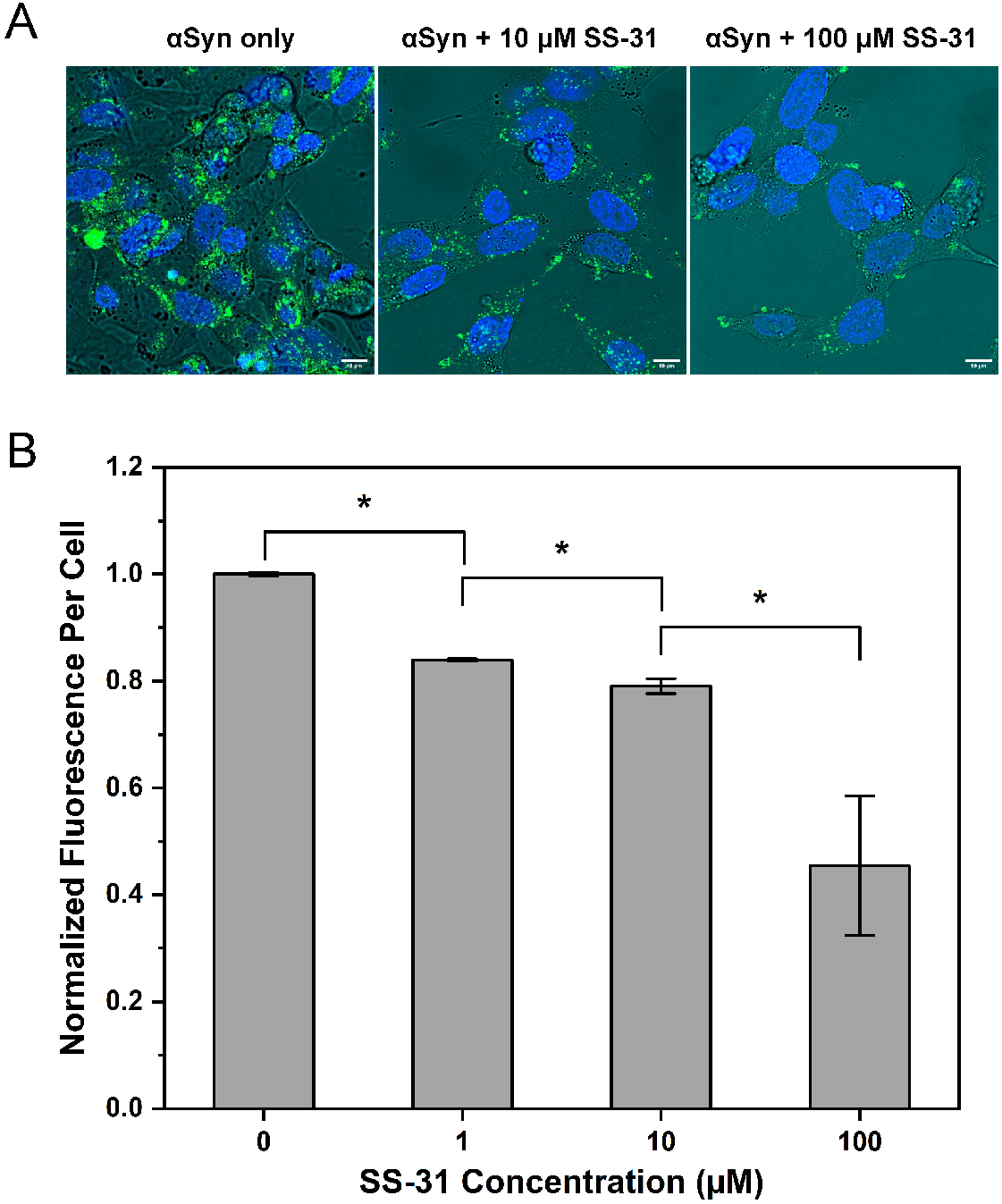
SS-31 affects cellular uptake of αSyn oligomers. (A) Confocal fluorescence images of BE(2)-M17 cells overlayed with the bright field images in the presence and absence of SS-31. The cell nucleus stained with Hoechst 33342 is shown in blue while the internalized αSyn is shown in green. (B) Comparison of the uptake of αSyn oligomers as indicated by the normalized fluorescence intensity per cell in different SS-31 concentrations. Statistical analysis was performed with student’s T-test (*: P<0.05).

## Discussion

Tetrapeptides composed of alternating cationic and aromatic residues, such as SS-31, are membrane and blood-brain barrier (BBB) permeable,^72^ an essential property for their potential to target brain diseases. In this study, we explored the promise of SS-31 as a modulator of lipid membrane binding and aggregation of αSyn. Using SUVs formed by DOPE/DOPS/DOPC (5:3:2 w/w) as a model lipid system, we observed distinct interactions between lipid membranes and both WT-αSyn and NAc-αSyn, revealing significant differences in their binding behaviors, in agreement with previous experimental and computational studies.^20,73^ The heightened affinity of NAc-αSyn to membrane can be attributed to the insertion of the terminal 14 amino acids of NAc-αSyn deeper into the membrane by approximately 2 Å.^20^ Importantly, our investigation extended to the impact of SS-31 on these interactions. Notably, NAc-αSyn exhibits greater resilience against displacement from lipid membranes by SS-31 than WT-αSyn, indicating that the replacement process may be also influenced by acetylation. These findings are crucial as the interaction between αSyn and cell membranes is pivotal for both its physiological function and aggregation process. Interfering with this interaction holds the potential to prevent the root culprit of DLB – the formation of toxic aggregates of αSyn.

Our study employed both fluorescence anisotropy and FCS techniques to unequivocally confirm that SS-31 effectively replaces αSyn from the membrane, of which FCS offers sensitivity at the level of fluorescence from single molecules. Moreover, extension at the cellular level, we revealed that SS-31 impedes αSyn entry into neuroblastoma cells through competitive lipid binding, as evidenced by confocal imaging. Here the positive charges that cationic residues of SS-31 carry and the hydrophobic interaction between the aromatic rings of the peptide and acyl chains of the lipid membrane may both contribute to such blockade effect. Good affinity with the lipid membrane and possession of positive charges enables SS-31 to modulate the electrostatics of the cell membrane,^74^ thus hindering the interaction of αSyn monomers with the membranes. A recent elegant microfluidic assay indicated that αSyn oligomers have a much higher affinity with the DOPS vesicles than αSyn monomers, hence displacing monomeric αSyn from lipid membranes.^75^ It is likely that SS-31 can also compete with αSyn oligomers for membrane binding, though to a lesser extent. Such a protective role of SS-31 could potentially prevent the cellular transmission of toxic αSyn oligomers.

Intriguingly, prior research has indicated that membrane binding accelerates αSyn nucleation and fibrillation.^28^ By displacing αSyn from the membrane, SS-31 appears to delay the onset of aggregation as well as prolong the aggregation process, hence inhibiting αSyn fibrillization induced by lipid membranes. TEM analysis of aggregated samples further revealed the impact of SS-31 on fibril morphology, i.e., suppressing amorphous aggregate formation and the development of twisted thick fibrils. Such morphological change may be attributed to multiple factors, including charge neutralization and di-tyrosine crosslinking between αSyn and SS-31.^76^

Moving to the cellular context, we investigated the influence of SS-31 on SH-SY5Y cells, a basic-level cellular model for Parkinson’s disease. The Seahorse Mito Stress Test revealed that SS-31 mitigates the impact of toxic αSyn oligomers on mitochondrial bioenergetics by enhancing basal respiration and maximal respiration, two parameters reflecting mitochondrial robustness and health, both were hampered by αSyn oligomers. Such mitigation can arise from several beneficial effects of SS-31. First, SS-31 acts by replacing αSyn in cytomembranes and inhibiting its aggregation; secondly, SS-31 scavenges reactive oxygen species by its tyrosine residue;^45^ and thirdly, SS-31 is concentrated in the inner mitochondrial membrane by interacting with cardiolipin, facilitating its interactions with components of the OXPHOS pathway, thus restoring OXPHOS complexes in mitochondria.^50^

In conclusion, our results presented here strongly suggest that alternating cationic and aromatic tetrapeptides, such as SS-31, are potential therapeutics for inhibiting α-Syn aggregation and alleviating mitochondrial stress, two hallmarks of Parkinson’s disease. It has been recently demonstrated that side chain composition and position influence the activity of these mitochondria-targeting peptides.^74^ Therefore, we envisage that there are great opportunities ahead for the rational design and covalent modification of such tetrapeptides with enhanced potency against many diseases linked to mitochondrial dysfunction and oxidative stress.

## Acknowledgements

This work was supported by the Leverhulme Trust (RPG-2015-345) and the Biotechnology and Biosciences Research Council (BB/R022429/1) of the United Kingdom. ES thanks the support from the European Union’s Horizon 2020 research and innovation program under the Marie Sklodowska-Curie grant agreement No 890595.

## Notes

### Competing Interest Statement

The authors have declared no competing interest.

