## Supplementary material for "Therapeutic Peptide SS-31 Modulates Membrane Binding and Aggregation of α-Synuclein and Restores Impaired Mitochondrial Function": Materials and methods, and supplementary figures

### Supplementary Information

Materials and Methods
Supplementary figures (Figure S1, Figure S2)
References

### Materials and Methods

#### Materials

SS-31 (Elamipretide) was purchased from Stratech Scientific Ltd, United Kingdom.

#### α-Synuclein Production and Purification

Wild-type α-synuclein (WT-αSyn) and N-terminal acetylated α-synuclein (NAc-αSyn) were expressed based on the protocol described previously.[1] The pT7-7 αSyn WT plasmid for WT-αSyn expression was obtained as a gift from the group of Professor Hilal Lashuel (Addgene plasmid # 36046 for WT-αSyn) and transformed into BL21(DE3) *E. Coli*. For NAc-αSyn expression, the BL21(DE3) *E. Coli* was transformed with both pT7-7 αSyn WT plasmid and pNatB (pACYC-duet-naa20-naa25) plasmid (Addgene plasmid # 53613). For the expression of protein, the recombinant cells were inoculated into 1L of LB medium containing 100 μg/mL ampicillin and incubated at 37 °C with 220 rpm shaking until the OD600 reached 0.7. Then, IPTG was added to a final concentration of 1 mM to induce αSyn expression. Subsequently, the cells were further incubated overnight at 28 °C with 220 rpm shaking and then pelleted by centrifugation at 8000 g at 4 °C. For cell lysis, the cell pellet was resuspended in Tris-HCl buffer (20 mM Tris-HCl, 50 mM NaCl, 5 mM EDTA, pH 7.4) with a protease inhibitor tablet (cOmplete, Roche) to prevent protein degradation. Then, the cell suspension was lysed by a sonic disembrator (Fisherbrand™ Model 120, 3×5 min, 25% cycles, 10% maximum power) with a replaceable probe (Fisherbrad™ 5/64 in., 2 mm). Afterwards, the cellular debris was spun down by centrifugation at 8000 g for 10 minutes. The supernatant was collected and filtered by 0.2 μm syringe filter to remove the cell debris. To precipitate nucleic acids, streptomycin sulfate was added to the supernatant to a final concentration of 10 mg/mL, and stirred for 15 minutes at -4 °C. After centrifugation at 16000 g for 20 minutes, the supernatant was collected, and ammonium sulfate was added to 50% saturation. The mixture was stirred for 30 minutes at 4 °C and centrifuged again at 16000 g. Finally, the pellet was collected, resuspended in Tris-HCl buffer (20 mM Tris-HCl,50 mM NaCl, pH 7.4) and dialysed overnight with 10 kDa dialysis membrane in 2L Tris-HCl buffer.

Crude protein after dialysis was purified by both 20 mL ion exchange column (HiPrep™ Q HP 16/10, Cytiva, GE Healthcare) and 320 mL size exclusion column (Hiloade™ 26/600 Superdex™ 75 pg, Cytiva, GE Healthcare) using the ÄKTA pure 25 L (Cytiva, GE Healthcare). For ion exchange column, samples were washed using Tris-HCl buffer (20mM Tris-HCl, 1 mM EDTA, pH 8.0), elution was performed using Tris-EDTA buffer (20 mM Tris-HCl, 1 mM EDTA, 1 M NaCl, pH 8.0). For size exclusion column, the sample was washed with water (MilliQ, Merck Millipore) and eluted by 1×PBS. Purified protein concentration was determined from the

UV absorbance at 280 nm using a molar absorption coefficient of 5960 $M^{-1}cm^{-1}$. Purity and acetylation of the protein were confirmed by electrospray ionization mass spectrometry (ESI-MS) using an orthogonal acceleration time-of-flight (oa-TOF) UPLC-MS system (WatersTM Aquity UPLC I-class, WatersTM LCT Premier Mass Spectrometer).

Single site mutated G7C αSyn was also produced as described previously.[1] The glutamine to cysteine at position 7 of αSyn was introduced using a Phusion Site-Directed Mutagenesis Kit (Thermo Fisher) following the manufacturer's protocol.

**Dye Labelling of α-Synuclein**

Alexa Fluor 488 C2 (Thermo Fisher Scientific) was used to label both WT-αSyn and NAc-αSyn that have a G7C mutation. Before labelling, a 10-fold molar excess of tris(2-carboxyethyl) phosphine (TCEP) was added to the protein solution to reduce disulphide bonds. After 30 minutes of stirring, the solution was eluted through a desalting column (Pd-10, GE Healthcare) to remove TCEP. Alexa 488 dye was then added to the protein sample in a molar ratio of 3:1. Subsequently, the mixture was stirred in the dark for 3 hours. Afterwards, the labelled protein was desalted using the desalting column and concentrated through a 10 K MWCO protein concentrator (Pierce, Thermo Fisher Scientific) to remove the excess dye. The purity of labelled protein samples was checked by SDS-PAGE, and the gel was imaged (IBright$^{TM}$ 1500 imaging System, Invitrogen) to ensure that free dye was negligible. Concentration of labelled protein was determined by Eq 1,

$$M = \frac{A_{280} - \gamma A_{495}}{\varepsilon_1} \qquad \text{(Equation 1)}$$

Where $A_{280}$ is the absorbance of the protein at 280 nm with a molar absorption coefficient $\varepsilon_1$ of 5960 $M^{-1}cm^{-1}$. $A_{495}$ is the absorbance of the dye at 495 nm with a molar absorption coefficient $\varepsilon_2$ of 72000 $M^{-1}cm^{-1}$, and $\gamma$ is a correction factor for fluorophore's contribution to the absorbance at 280 nm (0.11 for Alexa 488). The labelling efficiency $\eta$ can be calculated by Eq 2.

$$\eta = \frac{\varepsilon_1 A_{495}}{\varepsilon_2 (A_{280} - \gamma A_{495})} \qquad \text{(Equation 2)}$$

**Preparation of α-Synuclein Oligomer**

5 mL of WT-αSy in 50 mM HEPES (pH 7.4) and 0.02% $NaN_3$ was incubated at 37 °C for 20 hours without agitation. The excess αSyn monomer was removed by multiple filtrations using cut-off membrane (100 K MWCO protein concentrator, Pierce, Thermo Fisher Scientific). The oligomeric samples were used within the first two days after production. The concentration of

oligomers in monomer units was estimated based on the absorbance at 280 nm by using a molar extinction coefficient of 5960 $M^{-1}cm^{-1}$.

**Preparation of Lipid Vesicles**

All lipids were purchased as lyophilized powders from Sigma Aldrich, UK. They were dissolved in chloroform, then aliquoted and dried under a nitrogen stream overnight to obtain lipid films. To prepare small unilamellar vesicles (SUVs) with around 50 nm diameter, the lipid film of Coag Reagent I containing DOPE:DOPS:DOPC (5:3:2 w/w) was first hydrated in 50 mM HEPES buffer (pH 7.4) containing 100 mM NaCl with vortex for one hour. The lipid suspension was then extruded through a polycarbonate membrane with 50 nm pores (Nuclepore, Whatman) 19 times using an extruder (Mini-Extruder, Avanti Polar Lipids). The size of SUVs was confirmed by dynamic light scattering (DLS) using Zetasizer Ultra (Malvern Panalytical). The number of constituting lipids *n* can be estimated by Eq 3,

$$n = \frac{4\pi r^2 + 4\pi (r-l)^2}{a} \quad \text{(Equation 3)}$$

Where *r* is the radius of the vesicle, *l* is the thickness of bilayer and *a* is the average surface area per lipid. The number of constituting lipids in one 50 nm diameter SUV can be estimated to be ~$2.56\times10^4$ using 5.2 nm for *l* and ~ 0.5 $nm^2$ for *a*.

To make DMPS vesicles, 1,2-dimyristoyl-sn-glycero-3-phospho-L-serine (DMPS) lipid film was dissolved in n 20 mM phosphate buffer ($NaH_2PO_4$/$Na_2HPO_4$) (pH=6.5), 0.01% $NaN_3$ and stirred at 45°C for 2 hours. The resulting solution was then frozen and thawed five times under 45 °C water bath and dry ice. Then the obtained lipid suspension was sonicated (3 × 5 min, 50% cycles, 10% maximum power) on ice. The size of the DMPS vesicles was determined by dynamic light scattering (DLS) to be ~ 20 nm.

**Thioflavin T (ThT) Assay**

100 μM WT-αSyn or NAc-αSyn was incubated in 50 mM PBS buffer (pH 7.4) containing 0.01% $NaN_3$, in the presence of 50 μM ultrapure ThT, 100 μM DMPS and increasing concentrations of SS-31 (0 - 20 μM). CLARIOstar Plus plate reader (BMG Labtech Ltd, UK) was employed to monitor the fluorescence while incubating at 37 °C under quiescent conditions.

**Fluorescence Correlation Spectroscopy (FCS)**

FCS measurements were based on a custom-built confocal fluorescence setup based on an inverted optical microscope (Eclipse TE2000-U, Nikon) and equipped with a tuneable argon-ion laser (35LAP321-230, Melles Griot) as the excitation source. The fluorescence from the confocal volume passing through the confocal pinhole was split by a 50:50 beam splitter and detected by two detectors (SPCM-AQR-14 single photon counting module, PerkinElmer). Pseudo-auto correlation function was generated by a digital hardware correlator (Flex02-01D/C, Correlator.com). The setup used a 90 µm pinhole rendering it only sensitive to diffusion in the XY plane. Therefore, 2D diffusion models for one (Eq 4) and two species (Eq 5) were used to analyse the pseudo-autocorrelation curves.

$$G(\tau) = \frac{G(0)}{1+(\frac{\tau}{\tau_D})}\left(1 - F + Fe^{-\frac{\tau}{\tau_m}}\right) + C \quad \text{(Equation 4)}$$

$$G(\tau) = \left[\frac{G_1(0)}{1+\left(\frac{\tau}{\tau_1}\right)} + \frac{G_2(0)}{1+\left(\frac{\tau}{\tau_2}\right)}\right]\left(1 - F + Fe^{-\frac{\tau}{\tau_m}}\right) + C \quad \text{(Equation 5)}$$

The fractions of species $s_1$ and $s_2$ were simply estimated by Eq 6.

$$s_1 = \frac{G_1(0)}{G_1(0)+G_2(0)},\ s_2 = \frac{G_2(0)}{G_1(0)+G_2(0)} \quad \text{(Equation 6)}$$

**Fluorescence Anisotropy**

Fluorescence anisotropy measurements were performed using a spectrofluorometer (FluoroMax-4, Horiba). The binding between αSyn and SUVs was monitored by anisotropy while 100 nM labelled WT-αSyn or NAc-αSyn was titrated by various concentrations of the lipid. The binding affinity $k_d$ was determined by fitting fluorescence anisotropy $\theta$ to the Hill equation (Eq 7),

$$\theta = \frac{A}{1+K_d/[L]^n} + C \quad \text{(Equation 7)}$$

where *L* is the ligand concentration, *C* is the anisotropy value when *L* = 0, and *A* is the difference between the maximal anisotropy when binding is saturated and *C*. *n* is the Hill coefficient reflecting the binding cooperativity. The binding was assumed to be noncooperative, *n* = 1, in the data analysis.

The displacement of αSyn bound on SUVs by increasing SS-31 concentrations was monitored by fluorescence anisotropy in the same way. Anisotropy values were fitted by Eq 8, a sigmoid does-response function, to determine the concentration at half response, EC50,

$$\theta = A_1 + \frac{A_2 - A_1}{1+10^{(log\mathrm{EC50}-x)n}} \quad \text{(Equation 8)}$$

where x is the concentration of SS-31, $A_1$ and $A_2$ are the maximal and minimal anisotropy, respectively, $n$ is the Hill coefficient.

**Transmission Electron Microscopy (TEM)**

Negative staining transmission electron microscopy (TEM) was performed to analyse the morphology of samples after ThT assay. In short, 4 µL of samples were loaded onto an Agar carbon support grid, stained with 4 µL of 2% uranyl acetate then blot and left to dry. The thickness of the samples was less than 100 nm. Images were collected using a FEI Tecnai 12 120 kV BioTwin Spirit TEM in combination with a 2K Eagle CMOS camera and LAB6 filament source.

**Neuroblastoma Cell Culture**

Human SH-SY5Y cells were kept and grown at 37 °C in a humidified incubator chamber under an atmosphere of 5% $CO_2$ in Dulbecco's 100 modified Eagle medium (DMEM) supplemented with 10% (v/v) heat-inactivated fetal bovine serum (Life Technologies, UK). Cells were passaged 1-2 times per week, and the cells used for the experiments did not exceed 20 passages. The cells were plated when they reached 80-90% confluence. Human neuroblastoma BE(2)-M17 cells were cultured similarly and grown in Advanced Dulbecco's Modified Eagle Medium (DMEM) /Ham's F-12 medium supplemented with non-essential amino acids, 10% (v/v) heat-inactivated fetal bovine serum and 2 mM L-glutamine.

**Oxygen Consumption Rate Measurement and Mitochondrial Stress Test**

Bioenergetic profiling of neuroblastoma cells was performed using the Seahorse XFe96 Analyzer platform (Seahorse Bioscience, Agilent Technologies, UK). Oxygen consumption rate was measured during the assay, and bioenergetic parameters were acquired by using Seahorse XF Cell Mito Stress Test assay kit. Two days before the assay, 20,000 cells/well were seeded in 96-well plates, and experimental groups were exposed to 0 µM, 0.3 µM and 1 µM of wild-type αSyn oligomers. One day before the assay, new culture media (DMEM + 10% FBS) was changed, and the cells were exposed to 0 µM, 0.5 µM, 1 µM, 5 µM, and 10 µM of SS-31, on the same day, the XFe96 biosensor cartridge was activated using 1 ml of XF24 calibration buffer per well under 37 °C incubation overnight without $CO_2$. On the day of the analysis, cells were washed with Dulbecco's Phosphate-Buffered Saline (DPBS), and then the culture media was changed to 180 µL Seahorse DMEM Assay Medium, supplemented with 25 mM glucose, 2mM glutamine, and 2mM pyruvate, then, the cells were incubated at 37 °C in a humidified atmosphere of 5% $CO_2$ and 95% air for 1 hour. Cell Mitochondrial Stress Test was followed by basal respiration measurement. In brief, the mitochondrial effectors, oligomycin at 1 μM, carbonylcyanide-p-trifluoromethoxyphenylhydrazone (FCCP) at 2 μM, and a mixture of rotenone and antimycin A at 0.5 μM were injected sequentially and OCR was recorded in real-time. These reagents were included in Seahorse Mito Stress Test Kit, and the drug (SS-31) solutions were prepared by following the manufacturer's instructions. Data were analysed by the Seahorse Wave software.

**Confocal Imaging**

Confocal imaging was performed to visualise the intracellular distribution of Alexa 488 labelled WT-αSyn in BE(2)-M17 neuroblastoma cells. Cells were seeded with 30,000 cells/well in 8-well chambered coverglass (Thermo Fisher Scientific) and grown for at least 24 hours to reach 60 to 70% confluency before adding the labelled protein. Two days before the imaging, the culture medium was replaced with the same medium containing 100 nM Alexa 488 labelled WT-αSyn, and cells were exposed to αSyn for 24 hr in the incubator. One day before the imaging, various concentrations of SS-31 solution were added to different experimental groups. Cells were treated with SS-31 for another 24 hr before performing confocal imaging. Prior to imaging, cells were washed three times with PBS, stained by Hoechst 33342 (8 μM) for 10 mins, and washed again by PBS. Cells were then mounted for imaging using DPBS with $Ca^{2+}$ and $Mg^{2+}$. Imaging was carried out on a Leica STELLARIS 8 Inverted Confocal Microscope in the Facility for Imaging by Light Microscopy (Imperial College London) equipped with a HC PL APO 63x/1.40 OIL CS2 objective and a HyDS detector. All Images were analysed by FIJI. The corrected total cell fluorescence (CTCF) defined by Eq. 9 was calculated,

$$CTCF = IntDen - (Area \times Mean\ backgroud\ fluorescence\ readings) \quad \text{(Equation 9)}$$

where *IntDen* is the integrated intensity values, and *area* is the total area of each image. Mean fluorescence per cell determined by dividing CTCF by the cell number was then normalized to the control group (without SS-31).

**Statistical Analysis**

Statistical analysis was carried out in Origin 2024. Student's T-test was used and P-values of less than 0.05 were considered statistically significant.

### Supplementary Figures

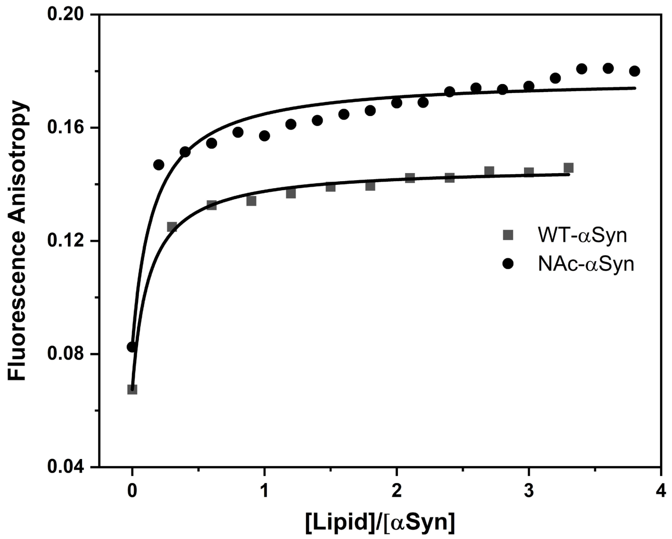


**Figure S1.** Fluorescence anisotropy titrations of wildtype and acetylated αSyn binding to synaptic-like SUV. Each data point was obtained by an average of 10 repeats. The measurements were performed in 50 mM HEPES at 298 K (pH 7.5).

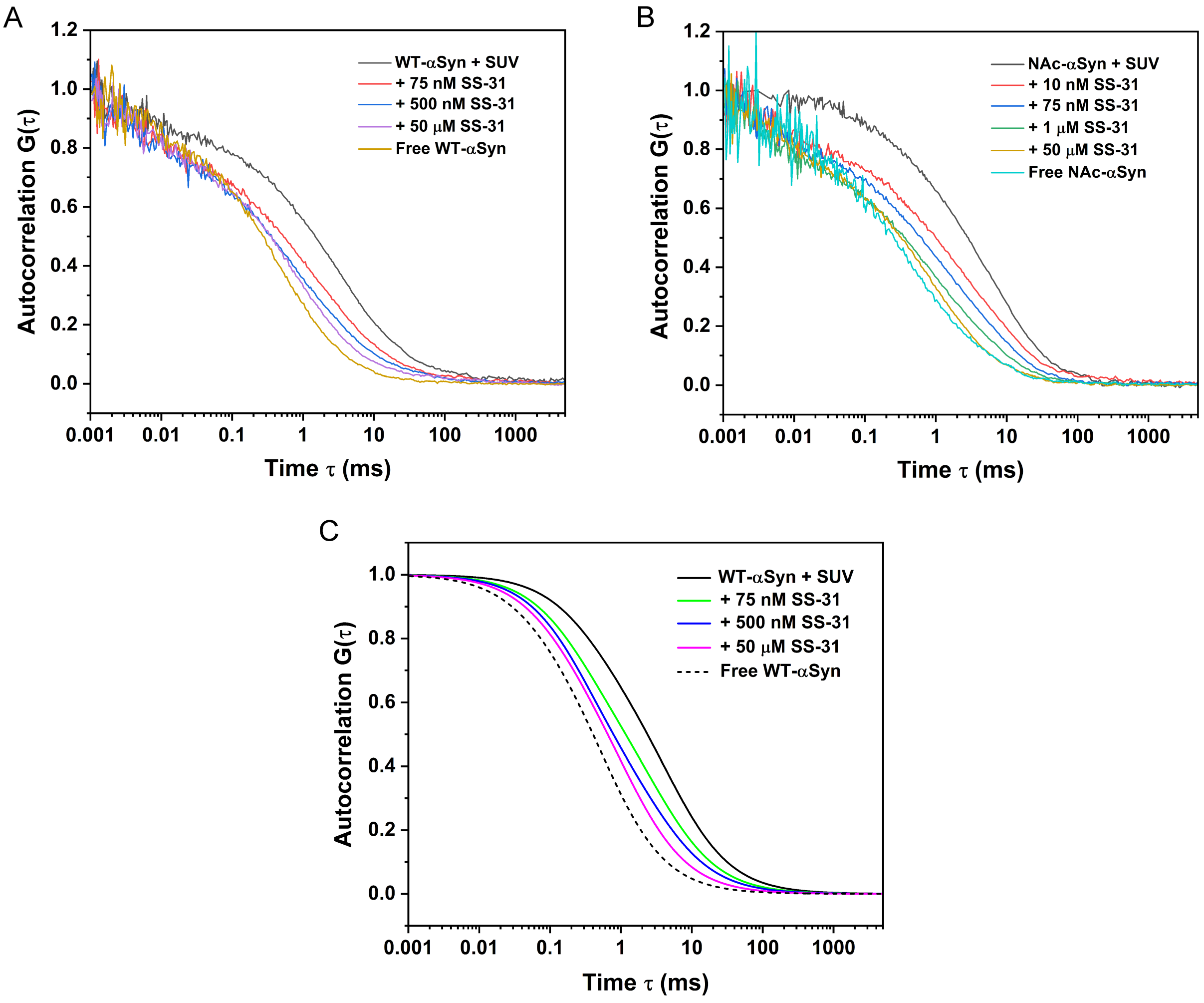


**Figure S2.** Normalized FCS curves for 100 nM labelled WT-αSyn (A) and NAc-αSyn (B) in the presence of 25 nM SUVs (0.5 mg/ml total lipid concentration) upon addition of various concentrations of SS-31. FCS measurements were performed in 50 mM HEPES solution at 298 K (pH 7.5). (C) Fitted diffusion only autocorrelation curves for the FCS curves shown in (A).
